# An intergenerational lipid memory of the social environment in *C. elegans*

**DOI:** 10.1101/2025.06.03.657568

**Authors:** Thomas Wilhelm, Wei-Wen Chen, Annika Wenzel, Wenyu Tang, Marcus T. Cicerone, Ben Lehner

**Affiliations:** Centre for Genomic Regulation (CRG), Barcelona Institute for Science and Technology (BIST), Barcelona, Spain; Wellcome Sanger Institute, Cambridge, UK; Universitat Pompeu Fabra (UPF), Barcelona, Spain; Institució Catalana de Recerca i Estudis Avançats (ICREA), Barcelona, Spain; School of Chemistry & Biochemistry, Georgia Institute of Technology, Atlanta, GA, USA; Neuro and Vascular Guidance Group, Buchmann Institute for Molecular Life Sciences (BMLS) and Institute of Cell Biology and Neuroscience, Frankfurt am Main, Germany

## Abstract

Information about the environment can, in some cases, be transmitted to an organism’s offspring by epigenetic inheritance. Here, we describe a novel form of epigenetics in *C. elegans* where information is transmitted between generations not by alterations in DNA, chromatin, or RNA, but by changes in the composition of lipids. Specifically, we delineate an environment-to-neuron-to-intestine-to-oocyte signalling axis that alters progeny thermotolerance by remodelling lipid provisioning to oocytes when animals detect social pheromones. Intergenerational information transmission via ‘lipid memories’ may represent an underappreciated form of epigenetics.

## Introduction

Neuroendocrine pathways enable the transmission of environmental information throughout an animal’s body, facilitating adaptation to changing conditions. The concept of the Weismann barrier, proposed by August Weismann in 1893, posits that such environmental information does not pass from somatic cells to germ cells ^1^. However, recent evidence challenges this view, showing that parental environments can sometimes influence offspring physiology, with information effectively crossing the Weismann barrier. This non-DNA sequence based ‘epigenetic’ transfer of information from parent to offspring can occur through multiple mechanisms, including changes in DNA methylation, altered histone modifications, and the provisioning of small RNAs ^2–4^.

Alternative routes of communication across the Weismann barrier that do not rely on these classical epigenetic mechanisms have also been described. For instance, in the nematode *C. elegans*, inter-generational memory of protein aggregation-induced neuronal mitochondrial stress has been proposed to be transmitted via elevated mitochondrial DNA levels ^5^. In addition, the inheritance of amyloid-like protein aggregates can affect multiple phenotypic traits, including germline development and sex differentiation ^6^. Beyond organelle- and amyloid-based intergenerational memory, we previously reported an example of intergenerational signalling where multiple differences between the progeny of older and younger nematodes are caused by altered provisioning of a lipoprotein complex to oocytes ^7^. Experimental manipulation of lipid composition in *C. elegans* further suggests a role of lipids in intergenerational signalling. For instance, feeding ursolic acid increases parental provisioning of sphingosine-1-phosphate to oocytes and thereby protects nerve cells from damage across two generations ^8^.

Here, by investigating variation in embryonic thermotolerance, we present an example of intergenerational signalling whereby environmental information is transferred between generations not by classic epigenetic mechanisms but by changes in the composition of lipids. We propose that such ‘lipid memories’ may be an underappreciated form of epigenetic inheritance.

## Results

### Maternal social environment modulates progeny thermotolerance

Embryonic development in *C. elegans* is generally extremely robust and predictable ^9,10^, yet under prolonged mild heat stress (28°C for 18 h), we observed that many worms arrest during late embryonic development. These embryos typically arrest at the 3-fold stage(Figure 1B), and do not hatch even after extended incubation (28°C for >30 h). The extent of this embryonic arrest varied across experiments, despite controlled conditions, prompting us to explore whether it was modulated by maternal environmental factors. Our previous work identified maternal age and pheromone sensing as two important influences on progeny phenotypes ^7,11^. We observed that offspring from densely populated plates exhibited increased heat resistance. Because maternal age and pheromone exposure could be confounded under these conditions, we investigated the effect of pheromone exposure while controlling for maternal age. To do this, we synchronised the parental generation at the L1 larval stage and introduced them to either worm-conditioned plates or untreated (control) plates. For hermaphrodite pre- conditioning, synchronised L1 larvae were placed on bacterial lawns and grown for three days, after which the worms and any remaining offspring were removed. For male pre- conditioning, synchronised *fog-2* males were grown on plates for five days and then removed, after which the parental generation was introduced (Figure 1A and S1A). The survival rate of embryos at high temperature was higher in the offspring of hermaphrodites exposed to plates pre-conditioned with hermaphrodites or males (Figures 1C and 1D). To rule out the influence of bacterial food availability or quality, we treated parental plates with either a filtered pheromone-containing extract from dense worm liquid cultures or a matched control extract from bacteria-only cultures (Figure 1A and S1A). Pheromone exposure improved offspring heat resistance similarly to plates pre-conditioned with worms (Figures 1E and 1F).

**Fig. 1.**
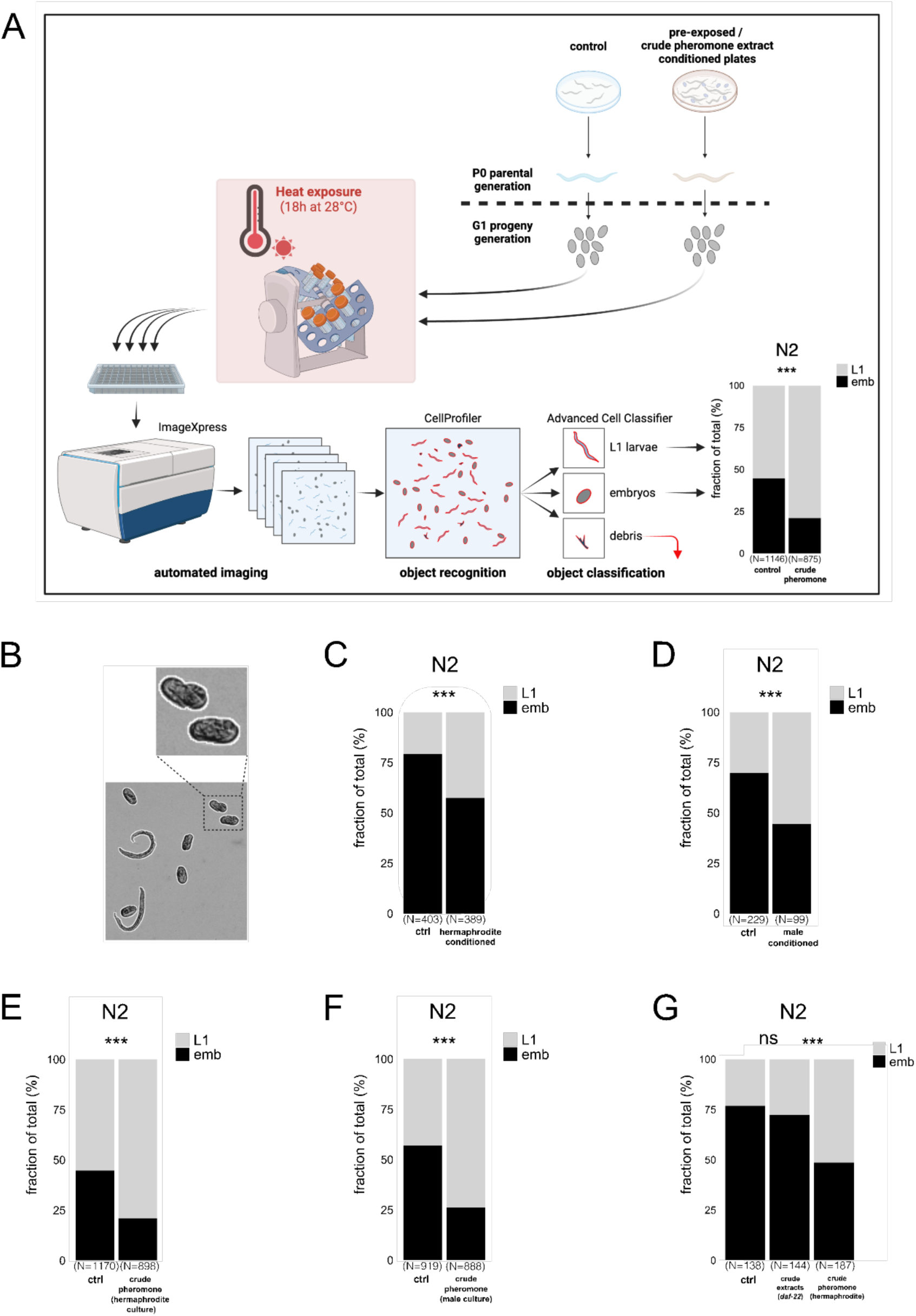
Maternal social environment modulates progeny heat resistance. **(A)** Schematic representation of the machine-learning-assisted screening platform. Parental (P0) generation nematodes were raised on plates pre-conditioned with crude pheromone extracts. Embryos from the offspring (G1) generation are separated post egg-lay and subjected to a prolonged mild heat stress (18h at 28°C). Hatching success was quantified using automated object recognition and classification. **(B)** Prolonged mild heat stress led to a consistent developmental arrest at the 3-fold stage in embryos that failed to hatch, as illustrated by a representative image. **(C, D)** Maternal exposure to secreted compounds via direct plate conditioning with hermaphrodites or males enhanced heat survival in the subsequent generation. **(E-G)** Maternal exposure to crude pheromone extracts from hermaphrodite or male cultures, but not from *daf-22* mutant cultures, significantly improved offspring survival under prolonged mild heat stress. **(A)** is created with BioRender.com. Statistical significance was assessed using Pearson’s Chi-squared test for independence **(C-G)**. ***P < 0.001; **P < 0.01; *P < 0.05; ns, not significant. P values and replicate experiments are provided in Table S1.

Ascarosides are a major class of *C. elegans* pheromones that regulate many aspects of physiology and behaviour ^12^. We prepared extracts from *daf-22* mutants, which are deficient in short-chain ascaroside production ^13^. These extracts failed to induce intergenerational heat-stress protection (Figure 1G).

### ASK chemosensory neurons and G protein signalling mediate intergenerational thermotolerance

Pheromones are detected by chemosensory neurons through G protein-coupled receptor (GPCR) signalling ^12^. *che-13(e1805)* mutants, which lack functional sensory cilia and cannot detect pheromones ^14^, failed to induce heat tolerance in their offspring after exposure to crude pheromone extracts (Figure 2A). *gpa-3* mutants, which lack a G-protein alpha subunit required for pheromone sensing and dauer entry in dense *C. elegans* cultures ^15^, also failed to induce offspring heat resistance after pheromone exposure (Figure 2B).

**Fig. 2.**
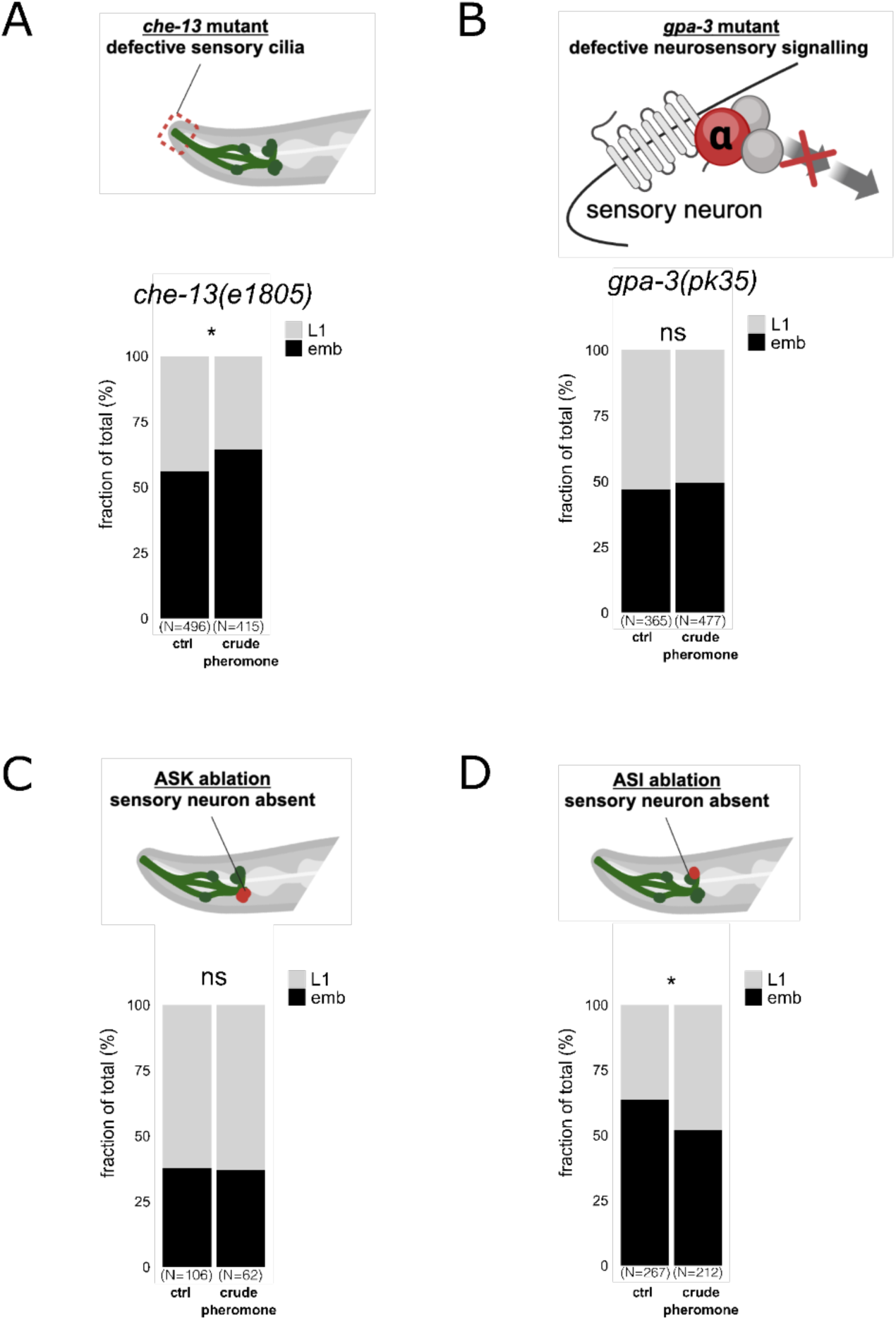
ASK chemosensory neurons and GPA-3 signalling are required for intergenerational heat tolerance. **(A)** The *che-13(e1805)* mutant worms, which lack functional chemosensory cilia, do not produce heat-stress-resistant offspring following maternal exposure to crude pheromone extracts. **(B)** Worms with a defective G protein α subunit (GPA-3) similarly fail to generate offspring with enhanced heat-stress resistance after maternal exposure to crude pheromone extracts. **(C, D)** Genetic ablation of ASK sensory neurons, but not ASI sensory neurons, inhibits the intergenerational increase in heat survival that follows maternal exposure to crude pheromone extracts. Elements of **(A-D)** are created with BioRender.com. Statistical significance was determined using Pearson’s Chi-squared test for independence (A-D). ***P < 0.001; **P < 0.01; *P < 0.05; ns, not significant. P values and replicate experiments are provided in Table S1.

Given that ASK neurons are essential for ascaroside pheromone-triggered behaviours and dauer formation ^16,17^, we next tested whether these neurons mediate intergenerational heat tolerance upon maternal pheromone exposure. Ablation of ASK neurons blocked the increased heat tolerance in offspring after maternal pheromone exposure (Figure 2C). In contrast, ablation of ASI neurons, which signal through DAF-7 to prevent dauer formation ^18^, did not impair the pheromone-induced heat tolerance in offspring (Figure 2D). Together, these results show that ASK neurons and GPA-3 signalling are required for pheromone-induced intergenerational heat tolerance.

### Maternal pheromone exposure triggers somatic and oocyte lipid remodelling

In *C. elegans*, pheromone sensing regulates lipid metabolism ^19^, leading us to hypothesise that changes in lipid provisioning might contribute to intergenerational heat resistance. Using GFP-tagged DHS-3, a lipid droplet protein ^20,21^, we observed a strong reduction in intestinal lipid droplet levels in hermaphrodites on male-conditioned plates (Figures 3A, 3B).

**Fig. 3.**
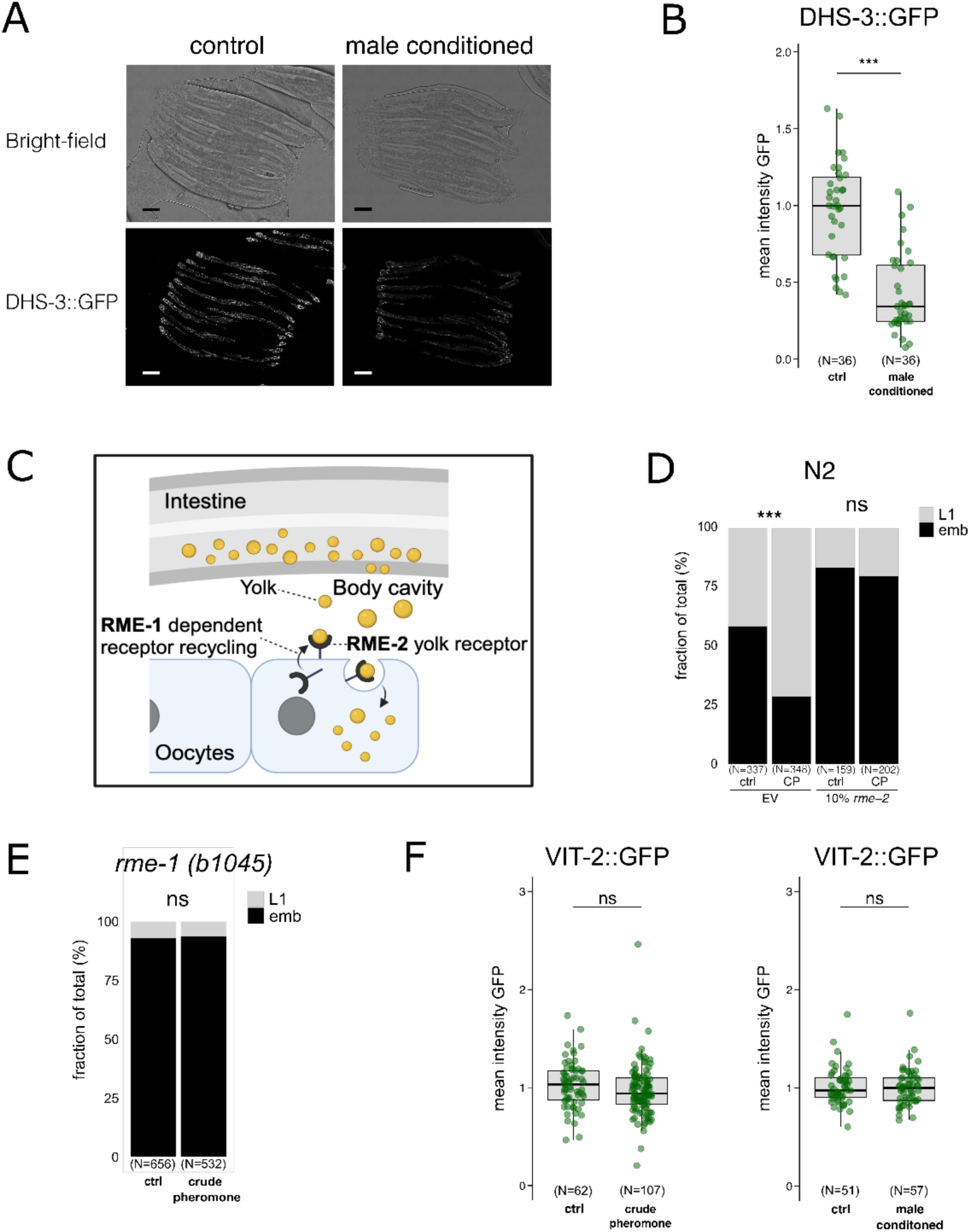
RME-2 receptor is required to transmit intergenerational heat resistance. **(A, B)** Worms grown on male-conditioned plates exhibit a reduction in intestine lipid droplets, quantified via DSH-3::GFP. Scale bar, 100 μm. **(C)** Schematic representation of RME-2-dependent yolk transport from the parent’s intestine to the oocyte. **(D)** Partial RNAi inhibition of the yolk receptor RME-2 diminishes intergenerational heat tolerance following maternal exposure to crude pheromone extracts. **(E)** *rme-1(b1045)* mutants, which are defective in yolk receptor recycling, similarly do not exhibit intergenerational heat resistance after exposure to crude pheromone extracts. **(F)** Boxplot depicting per-worm mean fluorescence of a yolk protein/vitellogenin fusion reporter, VIT-2::GFP, in 4- cell stage embryos from parents on control or pheromone-conditioned plates. Statistical significance was determined using the Mann-Whitney U test (B, F) or Pearson’s Chi-squared test for independence (D, E). **(C)** is created with BioRender.com. For all boxplots, the central line represents the median, and the box edges correspond to the first and third quartiles (25th and 75th percentiles), defining the interquartile range. ***P < 0.001; **P < 0.01; *P < 0.05; ns, not significant. Detailed P values and replicate experiments are provided in Table S1.

We next tested if parental lipid provisioning to oocytes contributes to the increased heat resistance in the offspring. Lipids are supplied to oocytes as lipoprotein yolk complexes that are secreted from the intestine and actively imported into oocytes via the RME-2 receptor ^22^ (Figure 3C). Inhibiting RME-2 expression using RNAi produced translucent eggs due to reduced yolk provisioning, and it did not severely affect egg-laying, consistent with previous observations ^7^. This reduction in RME-2 activity abolished the enhanced offspring survival at higher temperatures following exposure to crude pheromone (Figures 3D). Mutants lacking RME-1, an EH-domain protein required for RME-2 recycling and yolk uptake ^23^, also failed to produce offspring with increased heat resistance after maternal pheromone exposure (Figure 3C and 3E), further suggesting that lipoprotein uptake by oocytes is required for the response. However, we detected no increase in yolk, as quantified via the yolk protein VIT-2 after maternal exposure to crude pheromone or male-conditioned plates (Figure 3F). Hence, while the yolk uptake receptor is required to induce embryonic heat resistance, this is not associated with detectable changes in the amount of yolk provisioning.

The requirement for the RME-2 oocyte receptor but the lack of detectable changes in yolk provisioning led us to next test for pheromone-induced changes in maternal lipid content and yolk composition. We introduced two-photon fluorescence lifetime imaging microscopy (2p-FLIM) and broadband coherent anti-Stokes Raman scattering (BCARS) imaging for the further investigation. Using our recently established *in vivo* lipid-subtype quantifying approach with lipid dye Nile Red and 2p-FLIM ^24^, we first found a ∼50% reduction in total lipid in the maternal intestine following pheromone exposure (Figures S2A and S2B), corroborating our earlier observations of reduced levels of DHS-3 lipid droplets (Figures 3A and 3B). Secondly, taking advantage of the fact that the 2p-FLIM femtosecond laser can excite multiple dyes, we designed and performed a two-hour pulse feeding experiment to assess the fluorescent C12 fatty acid analog C1-C12-BODIPY signals in the intestine for studying lipid uptake, remodelling, and delivery. Our results revealed no significant difference in fatty acid uptake between pheromone-exposed and control animals (Figure S2D). However, pheromone-exposed hermaphrodites showed reduced triacylglycerol stores but increased polar lipids (Figures 4A, 4B, S2A, S2C). As the triacylglycerol-rich droplets act as somatic energy reserves and polar lipid-rich droplets resemble yolk composition ^25^, the reduction in intestinal lipid stores following pheromone exposure is likely driven by increased lipid mobilisation for yolk production rather than reduced uptake.

**Fig. 4.**
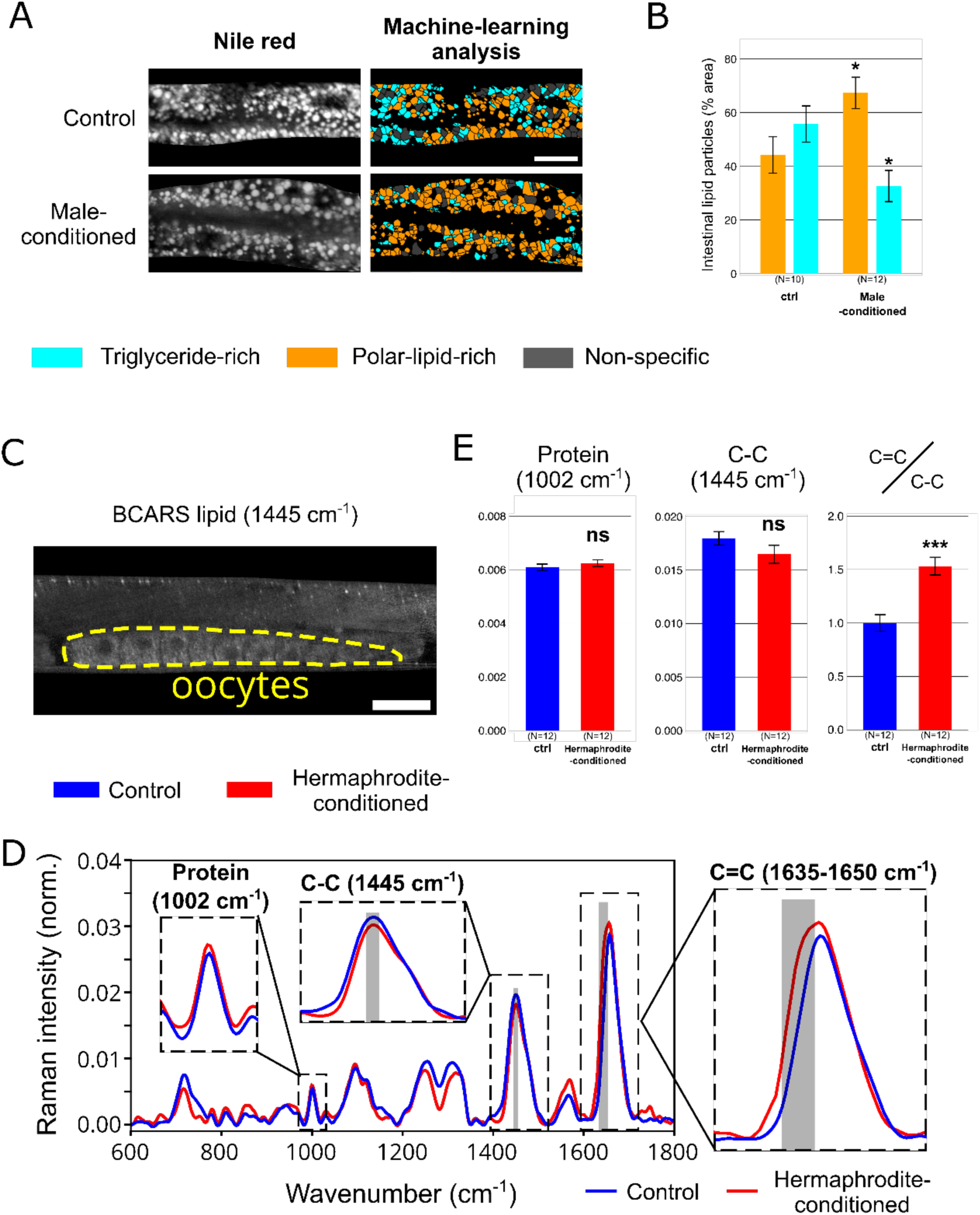
Maternal pheromone exposure causes somatic lipid remodelling and increases levels of unsaturated lipids in oocytes. **(A)** Left: Vital lipid staining through Nile Red 2p-FLIM in day 2 gravid wild-type hermaphrodites grown on male-conditioned or control plates. Right: Corresponding images displaying 2p-FLIM-based ensemble classification of triglyceride-rich (cyan) and polar-lipid-rich (orange) particles in the intestine. Scale bar,20 μm. **(B)** Bar chart quantifying 2p-FLIM-based ensemble lipid classification, distinguishing between triglyceride-rich (cyan) and polar-lipid-rich (orange) particles. **(C)** Lipid density Raman microscopy map at 1445 cm⁻¹ with a highlighted region of interest outlining oocytes within a day 2 gravid wild-type hermaphrodite, shown in yellow. Scale bar,30 μm**(D)** Raman spectra retrieved from oocytes of day 2 gravid wild-type hermaphrodites raised on hermaphrodite-conditioned plates (red line) versus control plates (blue line). The expanded frequency regions highlight the protein signal at 1002 cm⁻¹, saturated fatty acid carbon bonds at 1445 cm⁻¹, and unsaturated carbon bonds at 1635-1650 cm⁻¹. **(E)** Bar charts comparing the Raman spectra signals for protein (1002 cm⁻¹), saturated fatty acid carbon bonds (1445 cm⁻¹), and unsaturated carbon bonds (1635-1650 cm⁻¹) normalised to the fatty acid carbon bond signal. Data from oocytes of day 2 gravid wild-type hermaphrodites raised on hermaphrodite-conditioned plates (red bar) or control plates (blue bar). Bar charts represent mean values, with error bars indicating the SEM (B, E). Statistical significance was assessed using a two-tailed Student’s t-test (B, E). ***P < 0.001; **P < 0.01; *P < 0.05; ns, not significant.

Finally, we used BCARS imaging to characterise oocyte lipid composition ^25^ and assess whether maternal pheromone exposure alters the lipid make-up in the progeny. Since yolk lipoprotein uptake leads to delipidation ^25^, BCARS imaging provides a label-free method to detect intrinsic lipid composition without biases from lipid labelling efficiency of the oocytes in live hermaphrodites. Whilst we detected no significant changes in saturated fatty acids or protein content, maternal pheromone exposure led to a ∼50% increase in oocyte unsaturated fatty acids (Figures 4C-4E). Thus, in dense social environments with elevated pheromone exposure, maternal lipid metabolism undergoes a shift—marked by a significant depletion of intestinal lipid stores and an increased transfer of unsaturated fats to oocytes.

### A maternal lipid desaturation pathway confers intergenerational heat stress resistance

To determine how increased oocyte fatty acid unsaturation relates to parental desaturation activity, we comprehensively quantified the mRNA levels of all known C. elegans lipid desaturases in young adult hermaphrodites. Upon exposure to crude pheromone extract, mRNA levels for the Δ9 desaturase FAT-7, which, along with FAT-6 desaturates stearic acid (C18:0) to the monounsaturated fatty acid (MUFA) oleic acid (C18:1) ^26^, were two-fold increased (Figure 5A and 5B). mRNA levels for other desaturases involved in polyunsaturated fatty acid (PUFA) synthesis, such as FAT-1 and FAT-3 also showed modest but significant increases (Figure 5B). We confirmed FAT-7 upregulation using a *fat-7p::fat-7::GFP* reporter, detecting an increase in protein levels in L4 larvae, young adults, and adults following pheromone exposure (Figure 5C).

**Fig. 5.**
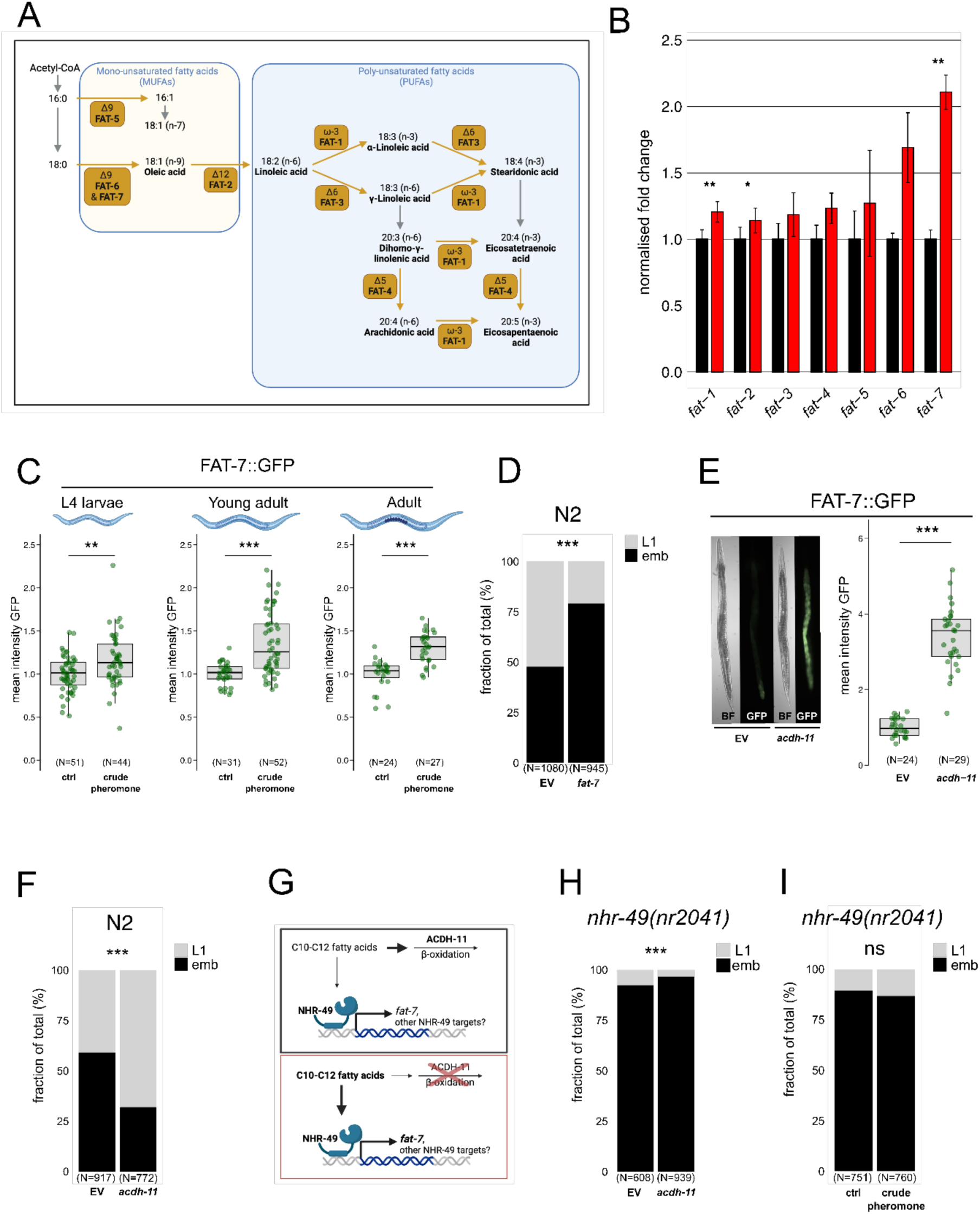
Maternal lipid desaturation confers intergenerational heat stress resistance. **(A)** Schematic diagram of fatty acid elongation and unsaturation pathways in *C. elegans*. **(B)** RT-qPCR analysis of fatty acid desaturase gene expression in young adults raised on crude pheromone-conditioned plates (red bars) versus control plates (black bars). Data represent fold changes relative to control across three independent biological replicates. Bar heights indicate mean fold change with error bars showing the SEM. **(C)** Boxplots depicting mean fluorescence of a *fat-7* fusion reporter, FAT-7::GFP, in L4 larvae, young adults, and day 2 gravid adults grown on crude pheromone-conditioned plates or control plates. **(D)** RNAi inhibition of *fat-7* via feeding dsRNA-expressing OP50 bacteria reduces offspring heat resistance under prolonged mild heat stress. **(E)** dsRNA-mediated RNAi inhibition of the medium-chain acyl-CoA dehydrogenase *acdh-11* improves offspring heat resistance under prolonged mild heat stress. **(F)** Quantification of FAT-7::GFP expression in day 2 gravid adults following dsRNA-mediated *acdh-11* RNAi inhibition compared to empty vector (EV) control. **(G)** Schematic summarising the suppression of *fat-7* expression by ACDH-11, which sequesters medium-chain fatty acids (C10-C12). In *acdh-11* mutants, *fat-7* is upregulated, a response previously attributed to the accumulation of medium-chain fatty acids that activate NHR-49-dependent *fat-7* transcription ^27,28^. **(H, I)** Neither *acdh-11* RNAi inhibition nor crude pheromone exposure enhances offspring heat stress resistance in *nhr-49(nr2041)* mutants. Statistical significance was determined using a paired two-tailed Student’s t-test (B), the Mann-Whitney U test (C, F), and Pearson’s Chi-squared test for independence (D, E, H, I). (A, G) and parts of (C) are created with BioRender.com. For all boxplots, the central line represents the median, and the box edges correspond to the first and third quartiles (25th and 75th percentiles), defining the interquartile range. ***P < 0.001; **P < 0.01; *P < 0.05; ns, not significant. P values and replicate experiments are provided in Table S1.

To test whether FAT-7 is required for progeny thermotolerance, we inhibited fat-7 expression using RNAi and quantified offspring survival at high temperatures. Offspring of *fat-7* RNAi-treated parents exhibited reduced heat resistance (Figure 5D). To genetically induce elevated fatty acid desaturation, we inhibited acyl-CoA dehydrogenase (*acdh-11*) through RNAi, a treatment known to upregulate *fat-7* expression ^27^. As expected, *acdh-11* inhibition strongly increased expression of the *fat-7p::fat-7::*GFP reporter (Figure 5E). This increase in *fat-7* expression correlated with a significant enhancement in offspring thermotolerance, as *acdh-11* RNAi-treated hermaphrodites produced heat-resistant progeny (Figure 5F). Previous studies showed that *acdh-11* inactivation enhances fat-7 expression via the nuclear hormone receptor NHR-49 ^27,28^ (Figure 5G). Consistent with this, *acdh-11* RNAi failed to enhance thermotolerance in *nhr- 49(nr2041)* mutant animals (Figure 5H). Furthermore, NHR-49 was required for pheromone-induced progeny thermotolerance, as *nhr-49* mutant worms failed to produce heat-resistant offspring following crude pheromone exposure (Figure 5I).

### Intestinal lipid remodelling induces offspring heat resistance

While other Δ9-desaturases such as FAT-5 and FAT-6 are expressed in various tissues, FAT-7 is exclusively expressed in the intestine ^29^. To inhibit gene expression specifically in the intestine, we used an intestine-specific RNAi strain *sid-1(q9);vha-6p::sid-1* (Figure 6A) ^30,31^. We confirmed that RNAi inhibition was confined to the parental intestine, as *pos- 1* RNAi, which normally causes germline-specific embryonic lethality, had no effect in *sid- 1(q9);vha-6p::sid-1* animals (Figure S3). Intestine-specific *acdh-11* inactivation increased offspring heat resistance (Figure 6B), with no further enhancement when combined with maternal pheromone exposure (Figure 6C). To further assess the role of FAT-7 in intergenerational heat resistance, we also reduced FAT-7 expression only in the hermaphrodite intestine and found that it reduced offspring survival at 28°C (Figure 6D).

**Fig. 6.**
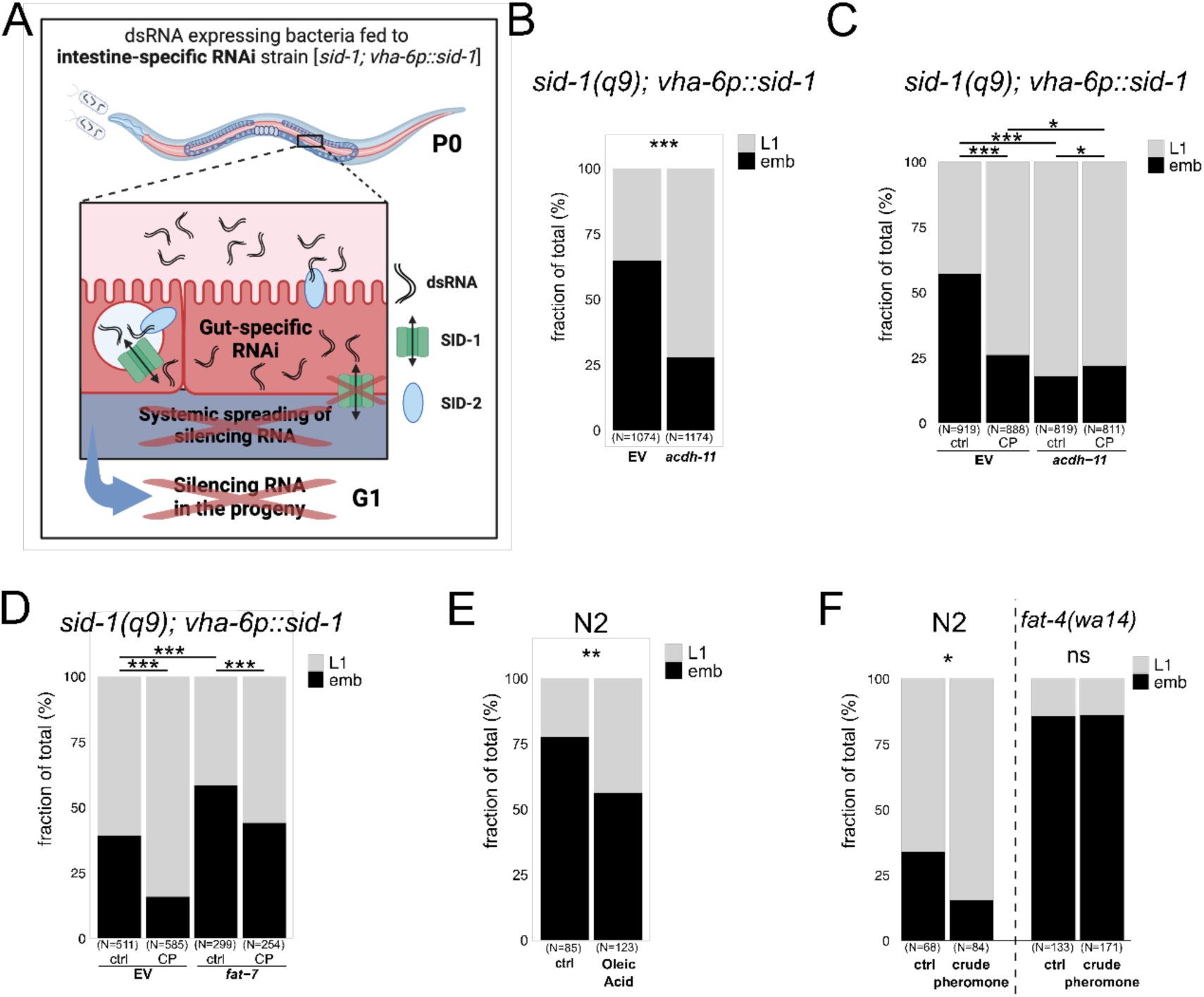
Intestine-specific lipid remodelling of fatty acid desaturation modulates offspring heat resistance. **(A)** Schematic of intestine-specific RNAi mutant. In wild-type *C. elegans*, ingested dsRNA is internalised via the SID-2 receptor and exported into the intestine cytoplasm by the SID-1 channel protein, which is also essential for dsRNA transport to other tissues. The intestine-specific RNAi mutant carries a mutation in *sid-1*, making it systemically RNAi-resistant, with *sid-1* expression selectively restored in the intestine using the *vha-6* promoter. Thus, ingested dsRNA triggers RNAi only in the intestine, preventing its spread to other tissues and the germline, blocking RNAi silencing across generations. **(B, C)** Maternal intestine-specific RNAi inhibition of *acdh-11* enhances offspring heat resistance. This effect does not further increase with maternal crude pheromone (CP) exposure. **(D)** Maternal intestine-specific RNAi inhibition of *fat-7* reduces offspring heat stress resistance. However, offspring from parents with intestine-specific *fat-7* RNAi inhibition still show improved survival under heat stress when exposed to crude pheromone. **(E)** Oleic acid supplementation (0.3mg per plate) in gravid adults enhances offspring resistance to heat stress. **(F)** *fat-4(wa14)* mutants produce offspring highly sensitive to prolonged mild heat stress. These mutants fail to mount intergenerational heat stress protection following maternal crude pheromone exposure. For this experiment, prolonged heat stress was applied at 27.5°C, as 28.0°C resulted in complete embryonic arrest in *fat-4(wa14)* mutants. **(A)** is created with BioRender.com. Statistical significance was assessed using Pearson’s Chi-squared test for independence **(B-F)**. ***P < 0.001; **P < 0.01; *P < 0.05; ns, not significant. P values and replicate experiments are provided in Table S1.

The Δ9-desaturases FAT-6 and FAT-7 convert stearic acid (C18:0) to oleic acid (C18:1), a cis-monounsaturated fatty acid (MUFA) ^32^. To examine oleic acid’s role, we tested *fat- 6;fat-7* double mutants, which lack Δ9-desaturase activity. These mutants displayed severe developmental delays at 20°C, making it impossible to assess their role during prolonged heat stress. However, feeding hermaphrodites with oleic acid enhanced offspring survival at 28°C, indicating its contribution to offspring heat resistance (Figure 6E). Oleic acid is the substrate for polyunsaturated fatty acid (PUFA) synthesis, involving its sequential desaturation and elongation (Figure 5A). We thus examined the role of PUFAs in pheromone-induced intergenerational heat resistance. The Δ12-desaturase FAT-2 initiates PUFA synthesis from oleic acid, and *fat-2* mutants, which largely lack all PUFAs ^33^, show severe growth and reproductive defects at 20°C. In contrast, *fat-4* mutants, lacking Δ5-desaturase activity and unable to produce long-chain PUFAs like arachidonic and eicosapentaenoic acids ^33^, developed normally at 20°C but were highly sensitive to mild prolonged heat stress (Figure 6F). Importantly, *fat-4* mutants also failed to produce heat-resistant offspring after maternal pheromone exposure, suggesting that specific PUFAs or their derivatives mediate the intergenerational heat resistance response.

In summary, intestine-specific fatty acid desaturation or direct oleic acid supplementation is sufficient to increase offspring heat resistance. Moreover, the enhanced heat resistance observed in offspring of pheromone-exposed mothers requires PUFA synthesis through FAT-4.

## Discussion

We have presented here evidence for an alternative mechanism of intergenerational epigenetic signalling in *C. elegans*, where environmental information is transmitted as a ‘lipid memory’ from parent to offspring to alter their phenotype and fitness (Figure 7A and 7B).

**Fig. 7.**
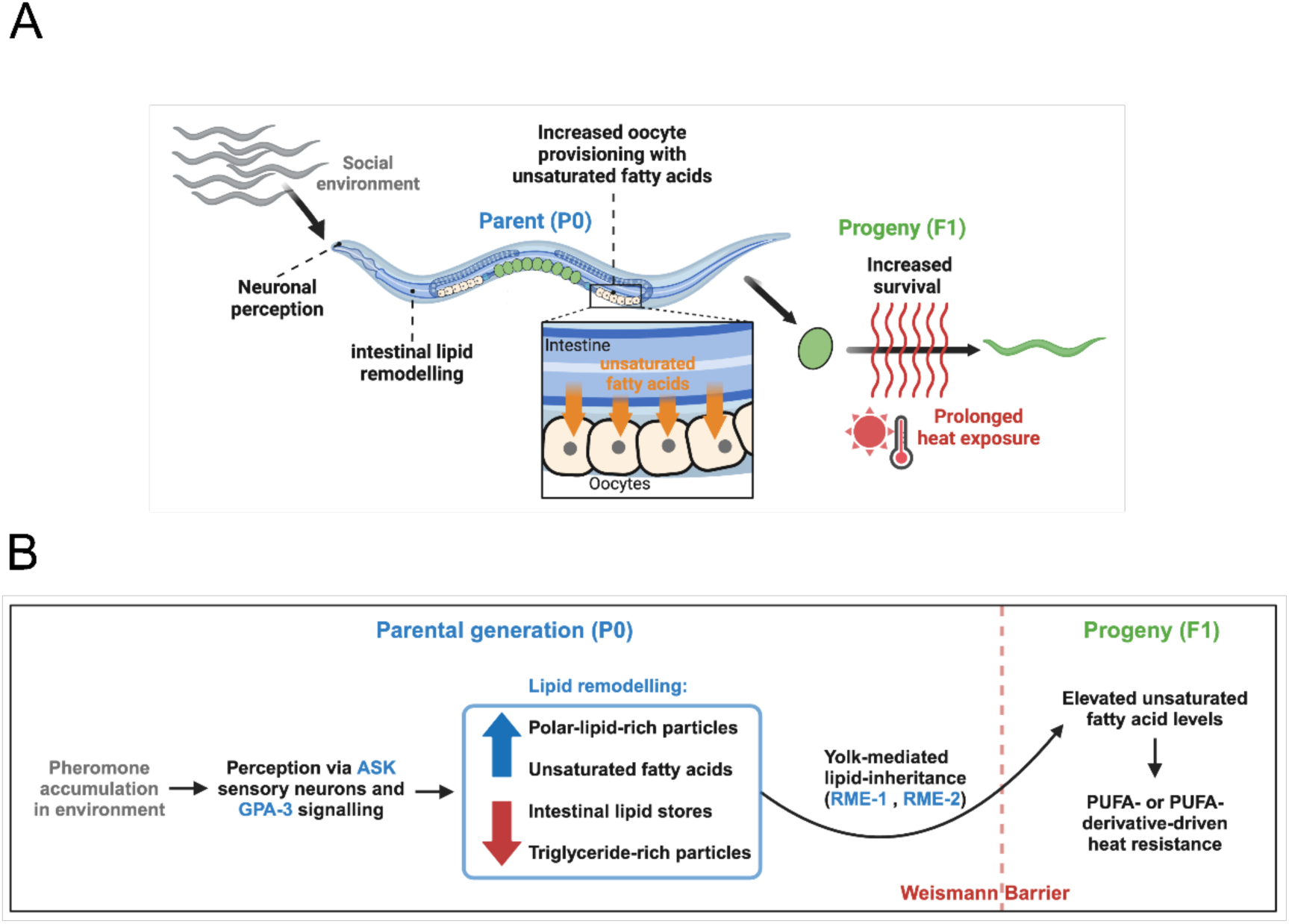
Intergenerational effects of the social environment in *C. elegans*. **(A, B)** Our findings demonstrate that population density, signalled through secreted compounds, modulates offspring resistance to prolonged mild heat stress (28°C for 18 h) in *C. elegans*. This environmental signal is detected by the ASK sensory neurons and requires GPA-3 signalling. Perception of high population density triggers extensive lipid remodelling, characterised by reduced intestine lipid stores and a relative increase in polar-rich lipids. We further show that yolk provisioning is crucial for the intergenerational transmission of heat stress resistance, with maternal intestine-specific modulation of lipid saturation playing a direct role in enhancing offspring fitness under elevated temperatures. Specifically, parents exposed to crude pheromone extract show increased provisioning of unsaturated fatty acids to their offspring. The essential role of the delta-5 desaturase FAT-4 in embryonic heat resistance, coupled with the dependency of the observed intergenerational phenotype on *fat-4*, suggests that the enhanced heat resistance in offspring of hermaphrodites experiencing higher population density operates through polyunsaturated fatty acids (PUFAs) or their derivatives. **(A, B)** are created with BioRender.com.

Our results show that chemosensory neurons, G-protein signalling, changes in intestinal lipid metabolism, and lipid uptake by oocytes are all required for this intergenerational response. Specifically, exposure to high pheromone levels induces intestine lipid remodelling, characterised by reduced triglyceride levels, increase in polar lipid levels, and upregulation of the Δ9-desaturase FAT-7. This increase in FAT-7 likely expands the pool of unsaturated fatty acids, including both the MUFA oleic acid and downstream PUFAs in the parental gut. Consistent with this, parents exposed to pheromones produce oocytes enriched in unsaturated fatty acids, and the Δ5-desaturase FAT-4 is also required for the response. In future work, it will be important to elucidate how this lipid memory protects embryos from heat stress.

In *C. elegans*, there is now evidence for at least three distinct lipid-mediated intergenerational signals. First, the total amount of vitellogenin yolk provisioned to oocytes controls larval size, developmental speed, and starvation resistance ^7^. Second, parental feeding with ursolic acid promotes intergenerational neuroprotection by enhancing the maternal provisioning of the signalling sphingolipid sphingosine-1-phosphate to oocytes^8^. Third, we have reported here that altered intestine lipid metabolism in response to neuronal perception of pheromones increases levels of oocyte unsaturated fatty acids, thereby enhancing embryo thermotolerance. Based on these results, we propose that, akin to how epigenetic memory transmits information across generations, shifts in maternally provisioned lipid composition may serve as a mechanism for intergenerational and transgenerational communication that is modulated by the ancestral environment. Future work should further test this hypothesis and explore both the diversity of lipid-mediated memories and the mechanisms by which they influence phenotypes and overall fitness across generations.

## Materials and Methods

### Nematode worms

Worms were cultured following standard procedures on nematode growth medium (NGM) plates as previously described ^34^, supplemented with 100 μg/mL streptomycin and 0.25 μg/mL Amphotericin B. Plates used were 90 mm in diameter with 26 mL of NGM unless otherwise specified. Worms were fed streptomycin-resistant Escherichia coli OP50-1 and maintained at 20°C unless stated otherwise. All nematode strains used in this study are listed in Table S1.

### RNAi

RNA interference (RNAi) treatments were performed using the RNAi feeding strain Escherichia coli OP50(xu363). RNAi plasmids from the Ahringer RNAi library were extracted from HT115 bacteria via miniprep, sequence-validated using the M13 forward primer, and transferred to OP50(xu363) through electrotransformation. For RNAi feeding, bacteria were grown overnight in LB medium, seeded at four times the native density onto NGM plates supplemented with 1 mM IPTG, 100 μg/mL ampicillin, and 0.25 μg/mL Amphotericin B. Plates were incubated overnight before introducing the worms to the bacterial lawn.

### Plate preconditioning with hermaphrodites or males

NGM plates were spotted in the centre with 700 μL of an overnight-grown OP50-1 culture and the lawn was grown one day at room temperature. For male conditioning, 40 synchronised *fog-2* L4-stage males were transferred onto the lawn with a platinum wire from a synchronised population. After five days, the worms were removed with a platinum wire. For hermaphrodite conditioning, 100-150 L1 larvae were spotted on the lawn and were washed off three days later. All laid eggs were allowed to hatch to L1 larvae before rinsing the plates at least five times with M9 buffer until no larvae were visible. Control plates were treated identically in parallel. After conditioning, the parental generation was introduced as L1 larvae at a density of 100-150 worms per plate. Plates showing any signs of contamination were excluded from the experiments. Neither the parental generation (P0) nor the progeny (G1) were starved at any point during the experiments.

### Crude pheromone preparation

Crude pheromone extract was prepared as described previously ^11^. Briefly, *C. elegans* were cultured in liquid NGM medium lacking peptone. After bleaching and hatching embryos overnight in M9 buffer to arrest at the L1 stage, larvae were transferred to a 250 mL NGM liquid culture in a 2.5 L flask at a density of four worms per μL. OP50-1 bacteria, grown overnight and washed twice with M9, were added at 50 mg/mL with 100 μg/mL streptomycin and 0.25 μg/mL Amphotericin B. Cultures were incubated at 20°C with shaking at 180 rpm for 4 days. Control cultures were grown in parallel with OP50-1 but without worms. Cultures were centrifuged at 200 g to remove worms, then the supernatant was centrifuged at 3000 g for 15 minutes to pellet bacteria. The clear supernatant was filtered through a 0.22 μm filter to remove any remaining bacteria, producing the crude pheromone, which was frozen in 1 mL aliquots at −70°C until use. 500 μL of the crude pheromone or control liquid was applied to the plates and allowed to dry completely (at least 6 hours) before spotting the parental generation as L1 larvae at a density of 100-150 worms per plate.

### Oleic acid supplementation

Oleic acid was supplemented as 18:1 (Δ9-Cis) PC (DOPC, Avanti Polar Lipids), dissolved in chloroform, and stored at −20°C. For each experiment, oleic acid (DOPC) was freshly dried under a nitrogen gas flow. The resulting lipid cake was further dried overnight (>16 h) in a vacuum chamber. The following morning, M9 buffer was added to each vial, and liposomes were formed by vortexing for 10 minutes at maximum speed. The negative control vial, containing only chloroform in the same volume as the lipid vial, was treated identically. Oleic acid (DOPC) liposomes, or the same volume of M9 for controls, were added to the worms on an OP50-1 lawn at a final concentration of 0.6 mg per 9 cm NGM plate and briefly dried under the laminar flow hood. After incubation at 20°C for about 7 hours, the offspring were harvested for the heat challenge (28°C for 18 h).

### Heat challenge of offspring embryos

82 to 86 hours after exposure to food as L1 larvae, the parental generation (P0) was washed off their plates as day two adults with M9 buffer, washed three times in 15 mL falcon tubes, and transferred to fresh OP50-1 plates. Eggs were laid within a 20–30- minute window, after which the parents were removed by washing the plates with M9 buffer (>5 times). Embryos remained on the plates and were collected by scraping with glass scrapers. Harvested embryos were washed 5 times with pre-heated (28°C) M9 buffer and placed in a 28°C incubator on a rotator inverting the tubes at 13 rpm for 18 hours. The next day, samples were spotted on 96-well glass bottom plates with sodium azide (NaN3) added at 20 mM to paralyse the worms for imaging. Automated imaging and progeny phenotype scoring followed (see Figure 1A and S1).

### Automated imaging and progeny phenotype scoring

Images were taken on 96-well glass bottom plates (MGB096-1-2-LG-L, Brooks) using an automated imaging system (ImageXpress Micro XLS, Molecular Devices) at 4x magnification. *C. elegans* objects were automatically identified using CellProfiler ^35^. Objects were classified as embryos, L1 larvae or debris via machine-learning-based identification with Advanced Cell Classifier (ACC) ^36^. Our ACC contained over 2600 objects (debris, embryos, and L1 larvae), achieving a classification accuracy of 97% (see Figure 1A and S1A for workflow).

### VIT-2::GFP quantification

Embryos from VIT-2::GFP reporter animals were isolated at day two of adulthood using standard hypochlorite treatment and collected in M9 buffer in an 8-well μ-slide (Ibidi) for imaging. Brightfield and epifluorescence microscopy were performed using a DMI6000B inverted microscope (Leica) equipped with an ORCA-Flash4.0 V2 Digital CMOS camera (Hamamatsu) at 20x magnification.

### DHS-3::GFP quantification

82 to 86 hours after exposure to food as L1 larvae, the parental generation (P0) was transferred to 2% agarose patches with 5 mM levamisole. Images were acquired immediately using a Leica SP8 STED confocal microscope at 20x magnification with fixed exposure parameters. Image analysis was performed in ImageJ ^37^ by measuring the area-corrected total cellular fluorescence (CTCF) for each worm ^38^. The intensity was calculated as follows: CTCF=((worm integrated density)-(background integrated density))/(worm area).

### Data visualisation and statistics

Statistical analyses were performed using R software ^39^. Data visualisation and figure generation were conducted using the ggplot2 package ^40^.

## Acknowledgments

This work was funded by a European Research Council (ERC) Advanced Grant (883742), the Spanish Ministry of Science and Innovation (LCF/PR/HR21/52410004, EMBL Partnership, Severo Ochoa Centre of Excellence), the Bettencourt Schueller Foundation, the AXA Research Fund, Agencia de Gestio d’Ajuts Universitaris i de Recerca (AGAUR, 2017 SGR 1322) and the CERCA Program/Generalitat de Catalunya. T.W. was funded by MSCA COFUND INTREPiD under the grant agreement No 754422. Some strains used in this study were provided by the Caenorhabditis Genetics Center, which is funded by NIH Office of Research and Infrastructure Programs (P40 OD10440). W.C and M.T.C. acknowledge support from NIH/NIA (R21AG086974). We thank all members of the Lehner lab for helpful discussions, and Marcos Francisco Perez and Antoni Beltran for providing valuable comments on the manuscript.

## Author contributions

T.W. conceived and performed experiments, analysed data, and drafted the manuscript. W.-W.C. and W.T. performed the experiments shown in Figure 4 and Figure S2. A.W. performed the experiments shown in Figure 3 A, B, and F. M.T.C. provided guidance and funding support for the experiments shown in Figure 4 and Figure S2. B.L. conceived experiments and edited the manuscript.

## Competing interests

The authors declare no competing interests.

## Data availability

All data are available in the main text or the extended data. Source data with P values and replicate experiments are provided in Table S1.

## Figures

**Fig. S1.**
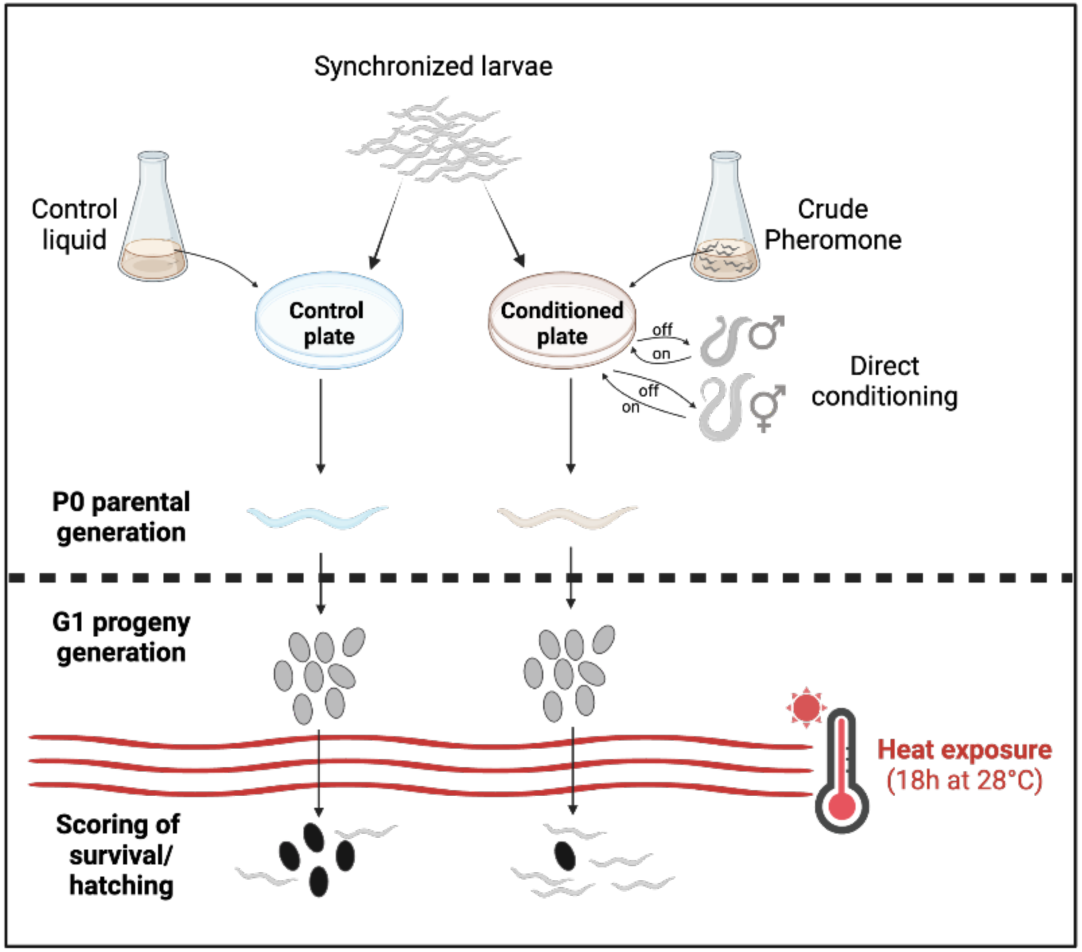
Maternal exposure to crude pheromone or male-conditioned plates does not affect in-utero developmental speed. Schematic representation of various plate conditioning methods that enhance intergenerational heat tolerance when the parental generation is exposed to these conditioned environments. Figure is created with BioRender.com.

**Fig. S2.**
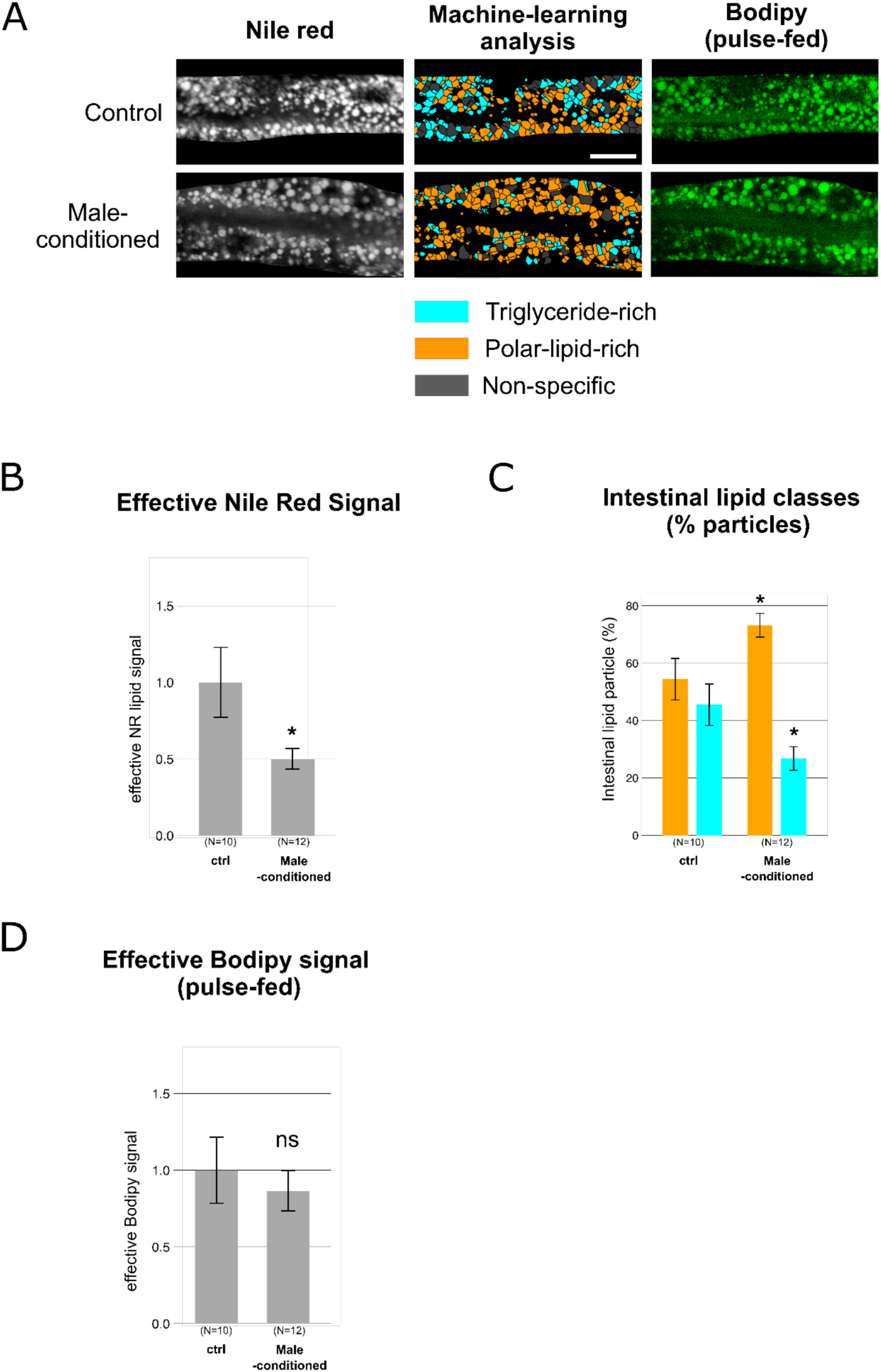
Male-conditioned plates reduce overall fat stores and cause lipid remodelling without significantly altering lipid uptake. **(A)** Left: Vital lipid staining using Nile Red 2p-FLIM in day 2 gravid wild-type hermaphrodites grown on male-conditioned or control plates. Middle: Corresponding 2p-FLIM images displaying ensemble classification of triglyceride-rich (cyan) and polar-lipid-rich (orange) particles in the intestine. Right: Signal from pulse-fed C1-BODIPY 500/512 C12 signal visualising lipid uptake. Scale bar, 20 μm. **(B)** Quantification of effective vital lipid staining using Nile Red 2p-FLIM in day 2 gravid wild-type hermaphrodites grown on male-conditioned or control plates. **(C)** Quantification of pulse-fed C1-BODIPY 500/512 C12 signal in day 2 gravid wild-type hermaphrodites grown on male-conditioned or control plates. Bar charts represent mean values, with error bars indicating the SEM. Statistical significance was determined using a two-tailed Student’s t-test (B, C). ***P < 0.001; **P < 0.01; *P < 0.05; ns, not significant.

**Fig. S3.**
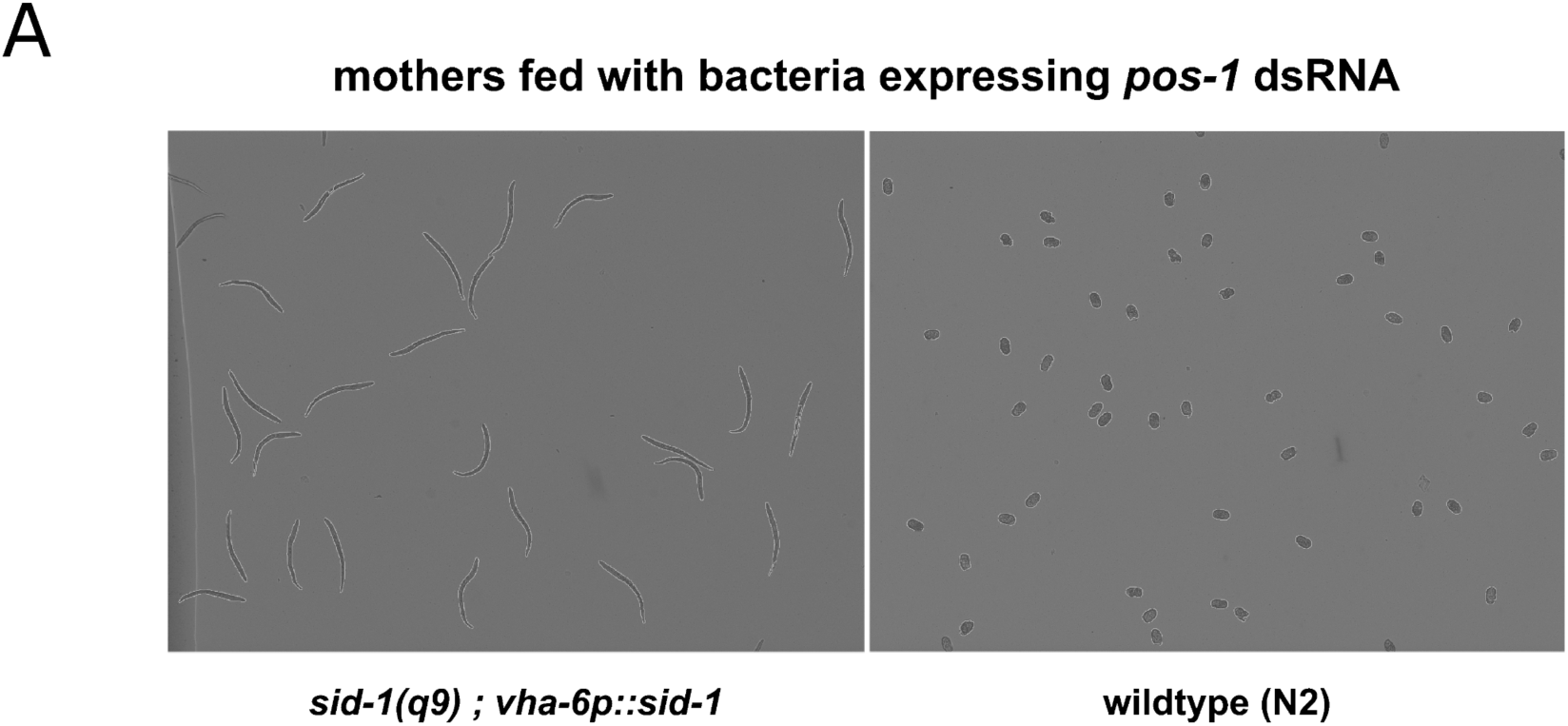
Maternal intestine-specific RNAi restricts RNAi effects to the intestine, preventing RNAi silencing in offspring. **(A)** Maternal intestine-specific *pos-1* RNAi via dsRNA feeding in *sid-1(q9); vha-6p::sid-1* mutants results in offspring unaffected by RNAi, while N2 wild-type worms exhibit complete embryonic arrest in their offspring due to early embryogenesis silencing. This indicates that maternal intestine-specific RNAi in *sid-1(q9); vha-6p::sid-1* mutants effectively prevents the spread of RNAi silencing to the germline and progeny. See also Figure 6A.

## Notes

### Competing Interest Statement

The authors have declared no competing interest.

